# Aging, matrix metalloproteinase imaging, and survival prospects in aortic aneurysm

**DOI:** 10.64898/2026.01.02.697375

**Authors:** Mean Ghim, Onur Varli, Azmi A. Ahmad, Afarin Neishabouri, Gunjan Kukreja, Zhengxing Zhang, Saalar Zarnegar, Nicholas Gangemi, Jakub Toczek, Jiasheng Zhang, Chi Liu, Yongjian Liu, Robert J. Gropler, Mehran M. Sadeghi

**Author notes:** Corresponding Author: Mehran M. Sadeghi, M.D., Professor of Medicine (Cardiology), Yale School of Medicine, 300 George Street, Room 770G, New Haven, CT 06511, USA.

## Abstract

Age is a risk factor for aortic aneurysm (AA), and different segments of the aorta exhibit varying susceptibilities to aneurysm. The specific factors that contribute to the higher incidence of AA and its complications with aging remain unclear. Matrix metalloproteinases (MMPs) are elevated in AA. However, the connection between aging, aortic MMP activity, and the increased prevalence of AA and its complications has not been systematically evaluated. This study leveraged MMP-targeted molecular imaging to investigate how aging affects aortic MMP expression and activity, as well as aneurysm development and survival.

**Methods:** AA development and animal survival were monitored for 28 days after Angiotensin (Ang)-II infusion in 8-10-week-old (young) and >51-week-old (old) *Apoe^−/−^* mice. Aortic MMP activation was quantified by PET/CT using an MMP-targeted tracer, ^64^Cu-RYM2, at baseline and 1 week after Ang II infusion. MMP activity and expression were quantified by tissue zymography and quantitative reverse transcription polymerase chain reaction, and compared between different segments of the aorta in young and old animals, and before and after Ang II infusion.

**Results:** Old animals’ survival to 28 days was significantly lower than that of young Ang-II-infused *Apoe^−/−^* mice (*P* < 0.05). ^64^Cu-RYM2 PET/CT showed significantly higher aortic MMP activation before and 1 week after Ang-II infusion in old compared to young *Apoe^−/−^*mice. The ^64^Cu-RYM2 signal was significantly higher in animals that did not survive 28 days than those that did (*P* < 0.01). MMP activity significantly increased by 4 days after Ang-II infusion, when dissection was found in a subset of *Apoe^−/−^* mice; and was significantly higher in the dissected, compared to adjacent, apparently normal, segments of the aorta. MMP activity was also significantly higher in the ascending thoracic aorta of untreated young and old mice, as well as of Ang-II-treated *Apoe^−/−^*mice (which was associated with significantly higher *Mmp2* gene expression), and of old wild-type mice.

**Conclusion:** Aging is associated with increased MMP activity along the aorta and worse AA survival. MMP-targeted molecular imaging can inform the aneurysm survival prospects. Selective MMP inhibitors and tracers may help prevent and track aneurysm growth, dissection, and rupture.

---

Each year, aortic aneurysms cause 15,000 deaths in the US and between 150,000 and 200,000 deaths worldwide (*1–3*). Advanced age is a major risk factor for developing abdominal aortic aneurysms (AAA). Notably, although AAA sizes display a similar age distribution among those screened for AAA, death from rupture is rare before age 65 (*4,5*). Advanced age is also a risk factor for thoracic aortic aneurysms (TAA) (*6,7*). TAA dissection in younger patients usually occurs in those with connective tissue disorders like Marfan and Ehlers-Danlos syndromes, but dissection of the more common “degenerative” TAA is more often seen in older patients (*7,8*). The combined effects of risk factors like hypertension and smoking increase with age, but the specific factors that contribute to or mediate the higher incidence of aortic aneurysms as age advances are still unclear. Different parts of the aorta contain smooth muscle cells (SMC) of various embryologic origins and unique biomechanical properties, which respond differently to aging and injury (*9,10*). As such, human AAA often involves the infra-renal aorta (IRA) (*4*) and a suprarenal aorta (SRA) predisposition is seen in an established model of aortic aneurysm, namely angiotensin II (Ang II)-induced aneurysm in apolipoprotein-deficient (*Apoe^−/−^*^)^ mice (*11*).

Matrix metalloproteinases (MMPs) are elevated in aortic aneurysms and play a role in aneurysm development (*12*). In aortic aneurysm tissues from patients undergoing surgery for aneurysm complications, aortic dissection and rupture, MMP expression and activity are upregulated compared to the normal aorta (*12,13*). Similarly, MMPs are upregulated in animal models of aortic aneurysms (*12*). While it is likely that increased MMP expression and activity contribute to the pathogenesis of aneurysm complications, one cannot rule out that the increased MMP activity observed in such surgical specimens results from dissection or rupture that prompts surgical intervention. The expression of some members of the MMP family is increased with aging (*14,15*). However, the relation between aging and MMP activity, and its contribution to the increased prevalence of aortic aneurysms and their complications with aging, has not been systematically evaluated.

MMP-targeted molecular imaging is a promising approach for tracking vessel wall biology in vivo and may be valuable for aneurysm risk stratification. Recently, several positron emission tomography (PET) and single photon emission computed tomography (SPECT) MMP-targeting radiotracers, such as ^64^Cu-RYM2 and ^99m^Tc-RYM1, have been developed and shown promise for detecting aortic aneurysms in murine models (*13,16,17*). Hypothesizing that aging is associated with increased aortic MMP activity and predisposes the aorta to aneurysmogenic stimuli, here we leveraged MMP-targeted molecular imaging in combination with tissue zymography and gene expression analysis to a) assess how aging affects aneurysm development and survival, b) investigate the effect of aging on aortic MMP expression and activity, c) evaluate the performance of MMP-targeted molecular imaging (^64^Cu-RYM2 PET/CT) in predicting aneurysm outcome, and d) elucidate the spatial and temporal relationship between MMP activity and aortic dissection.

## Materials and Methods

Detailed materials and methods are provided in the supplemental appendix.

### Animals

Animal experiments were performed under protocols approved by the Institutional Animal Use and Care Committees of Yale University and the Veterans Affairs Connecticut Healthcare System. Male C57BL/6J (wild-type, WT, n = 11) and apolipoprotein E-deficient (*Apoe^−/−^*, n = 106) mice, originally from Jackson Laboratory, were bred in-house. A flow chart summarizing animal experimental groups is provided in the Supplemental Material (Supplemental Figure 1). In vivo imaging studies focused on 8-10 weeks old (“young”) and >51 week-old (“old”) mice, while tissue analysis included an additional intermediate age, fully adult (12-17 week old) group of animals. To induce aortic aneurysm, *Apoe^−/−^* mice were infused with recombinant human Ang II (Santa Cruz Biotechnology) at 1,000 ng/kg/min for up to 28 days via osmotic minipumps (models 2001 or 2004; Alzet) implanted subcutaneously under isoflurane anesthesia. Postoperative analgesia was provided as a single subcutaneous dose of extended-release buprenorphine (Ethiqa XR, 3.25 mg/kg).

### RYM2 labeling

Radiolabeling of RYM2 with ^64^Cu was performed following previously reported procedures (*13*). The specific activity of ^64^Cu-RYM2 was 78.2 ± 8.4 GBq/µmol.

### PET/CT imaging

^64^Cu-RYM2 PET/CT was performed as described (*13*). Briefly, under inhaled isoflurane anesthesia, the animals were injected intravenously with 11.73 ± 3.5 MBq of ^64^Cu-RYM2 and 30 µL + (body weight in grams) µL of Exitron nano 12000 (Viscover Imaging). A CT acquisition was then carried out, followed by a 10-minute PET acquisition on a small-animal dedicated PET/CT scanner (Inveon PET/CT, Siemens Medical Solutions), beginning 50 minutes after tracer injection.

### Statistical analysis

Data are presented as mean ± standard deviation (SD). Normality of the data distributions was assessed using the Shapiro-Wilk test before applying parametric statistical tests. Unpaired or paired two-tailed t*-*tests were used to compare two groups. For comparisons involving more than two groups, one-way analysis of variance (ANOVA) with Tukey’s post hoc multiple-comparison test was used. Repeated-measures (RM) ANOVA was used for matched measurements. Kaplan–Meier survival curves were compared using the log-rank (Mantel–Cox) test. A *P* value < 0.05 was considered statistically significant.

## Results

### Animal age and survival in aortic aneurysm

Angiotensin II infusion triggers aortic aneurysm development and death from rupture in *Apoe^−/−^* mice. A comparison of animal survival to 28 days in young (8-10 weeks, n = 18) and old (>51 weeks, n = 11) *Apoe^−/−^* mice infused with Ang II showed significantly less survival in older animals (Figure 1A, *P* < 0.05). Morphometric analysis of the aorta in surviving animals showed the presence of AAA in 5 out of 14 young and 1 out of 4 old mice (Figure 1B).

**Figure 1.**
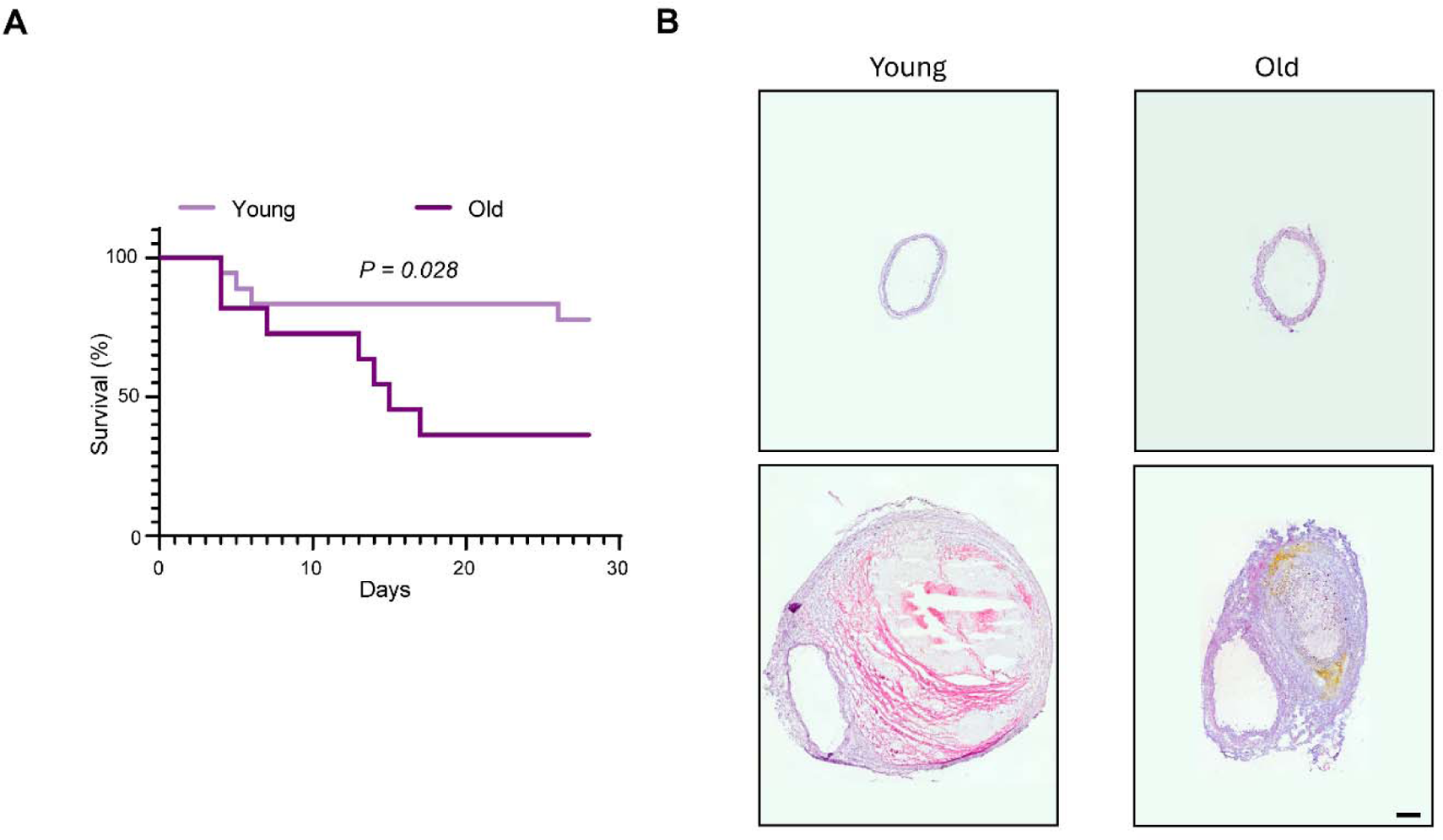
Age and survival in angiotensin II-infused *Apoe^−/−^* mice. **A.** Kaplan–Meier survival curves for young (8–10 weeks) and old (>51 weeks) *Apoe^−/−^* mice during 28 days of angiotensin II infusion (*P* < 0.05). **B.** Illustrative examples of H&E-stained cross-sections of the suprarenal abdominal aorta from young and old mice without (top panels) and with AAA (bottom panels). Scale bar = 200 μm.

### Animal age, aorta MMP activation, and aneurysm outcome

MMPs mediate aortic aneurysm development and rupture. In vivo PET/CT imaging using ^64^Cu-RYM2 (a tracer that targets activated MMPs (*13*)) on day 7 of Ang II infusion showed significantly higher aorta tracer uptake in old mice compared to young animals (descending thoracic aorta: SUVmax 0.96 ± 0.20 in young mice vs 1.36 ± 0.39 in old mice, *P* < 0.01; suprarenal abdominal aorta: SUVmax 1.17 ± 0.23 in young mice vs 1.80 ± 0.52 in old mice, *P* < 0.01, Figure 2). Notably, tracer uptake in the suprarenal abdominal aorta was significantly higher in animals that did not survive 28 days than those that not (SUVmax 1.21 ± 0.25 in surviving animals vs 2.11 ± 0.45 in non-surviving animals, *P* < 0.0001, Figure 3 and Supplemental Figure 2).

**Figure 2.**
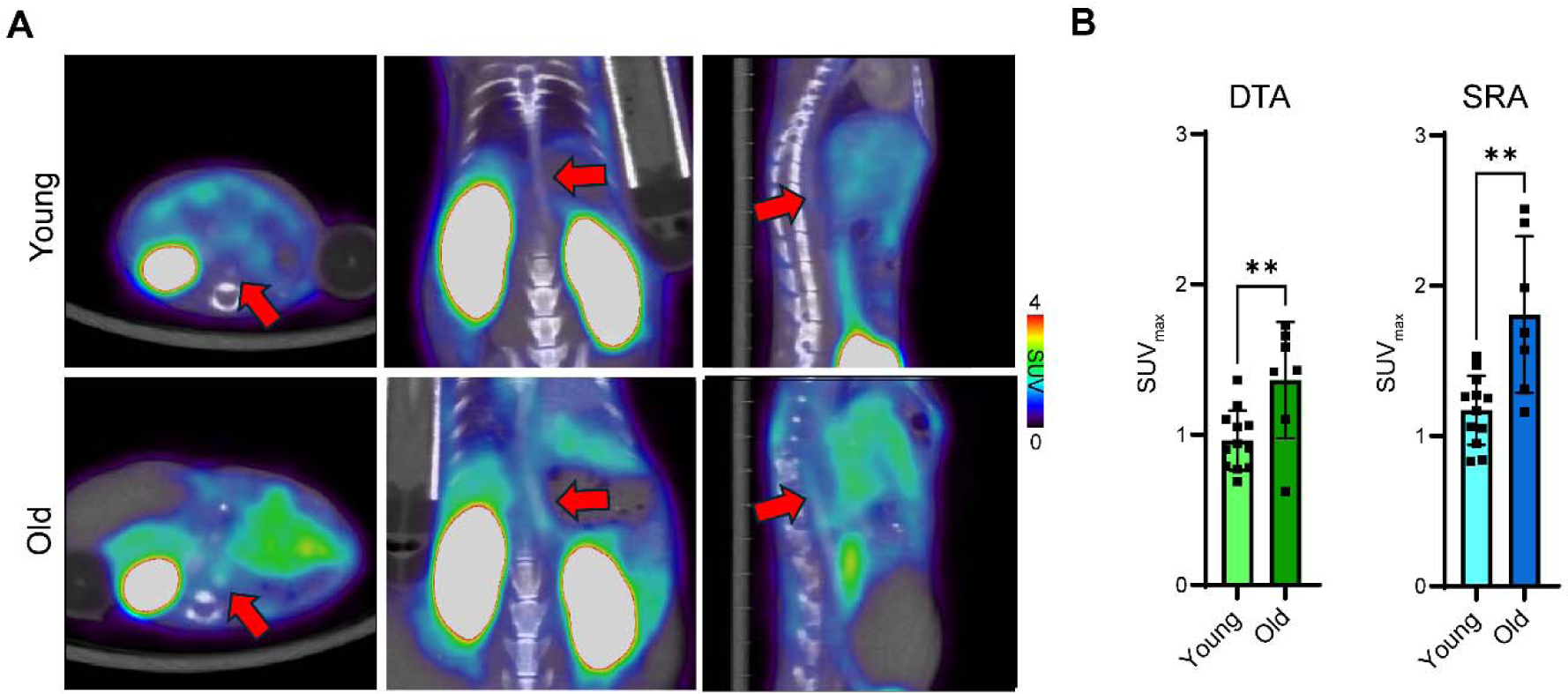
Age and ^64^Cu-RYM2 PET/CT in angiotensin II-infused *Apoe^−/−^* mice. **A.** Illustrative examples of fused ^64^Cu-RYM2 PET/contrast-enhanced CT images (axial, coronal, and sagittal) acquired 7 days after starting angiotensin II infusion in young (8–10 weeks) and old (>51 weeks) *Apoe^−/−^* mice. Red arrows indicate the suprarenal abdominal aorta. **B.** Quantification of the ^64^Cu-RYM2 signal in the descending thoracic aorta (DTA) and suprarenal abdominal aorta (SRA) in young and old mice. **: *P* < 0.01. SUV: Standardized uptake value.

**Figure 3.**
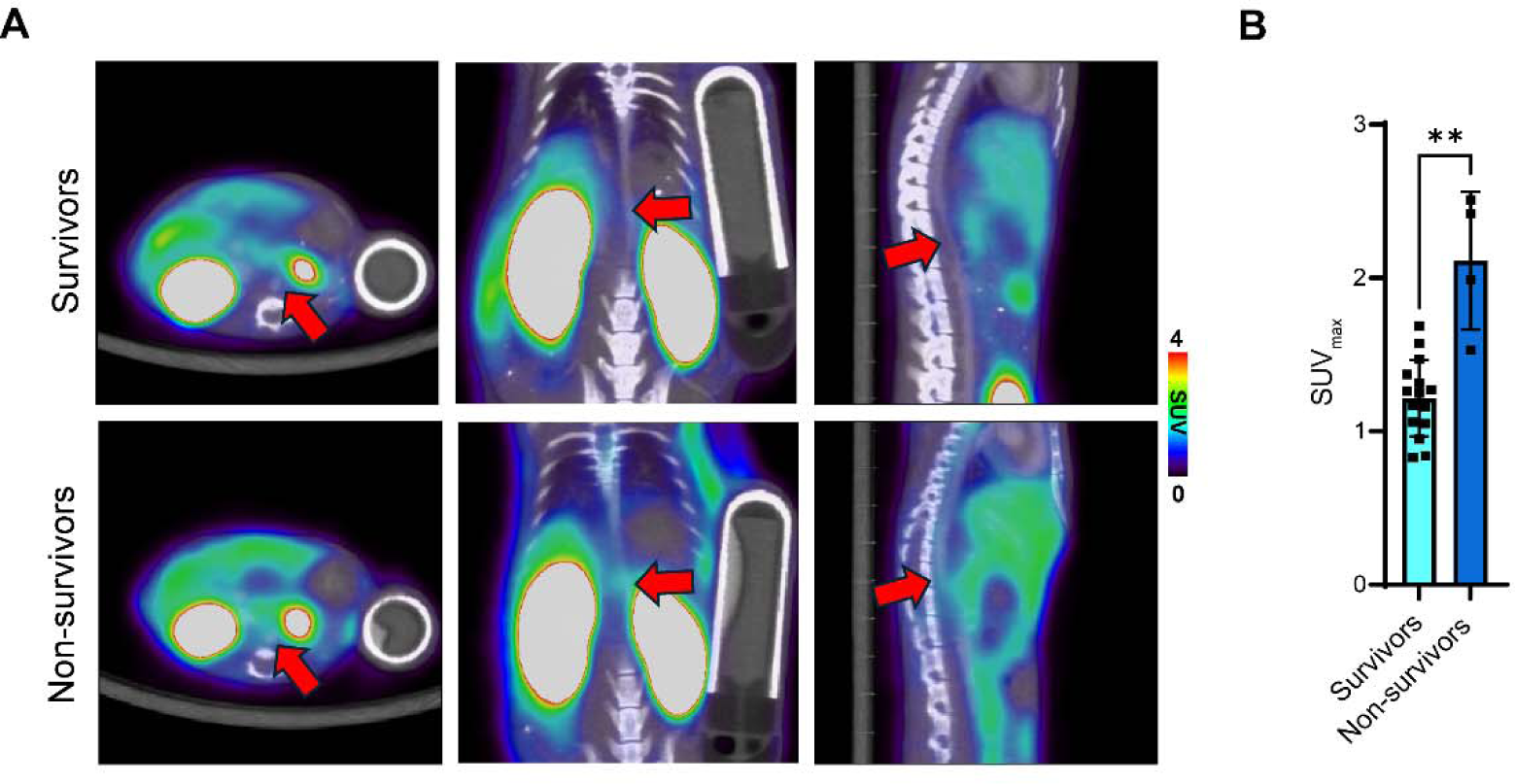
^64^Cu-RYM2 PET/CT and survival in angiotensin II-infused *Apoe^−/−^* mice. **A.** Illustrative examples of fused ^64^Cu-RYM2 PET/contrast-enhanced CT images (axial, coronal, and sagittal) acquired 7 days after starting angiotensin II infusion in young (8–10 weeks) mice that survived to day 28 (survivors) and from mice that died before day 28 (non-survivors). Red arrows indicate the suprarenal abdominal aorta. **B.** Quantification of the suprarenal abdominal aorta ^64^Cu-RYM2 signal on PET/CT images acquired on day 7 in young and old non-survivors and survivors. **: *P* < 0.01. SUV: Standardized uptake value.

To investigate whether the higher aorta MMP activation in Ang II-infused old mice compared to young animals is also present without Ang II infusion, a group of young and old *Apoe^−/−^* mice underwent ^64^Cu-RYM2 PET/CT imaging. This showed significantly higher suprarenal abdominal aorta tracer uptake in old mice compared to young animals (SUVmax 0.60 ± 0.05 in young mice, vs 0.84 ± 0.07 in old mice, *P* < 0.001, Figure 4). In both young and old animals, the suprarenal abdominal aorta MMP signal on ^64^Cu-RYM2 PET-CT images was significantly higher after 7 days of Ang II infusion compared to untreated mice (*P* < 0.0001 for young and *P* < 0.01 for old animals, Supplemental Figure 3).

**Figure 4.**
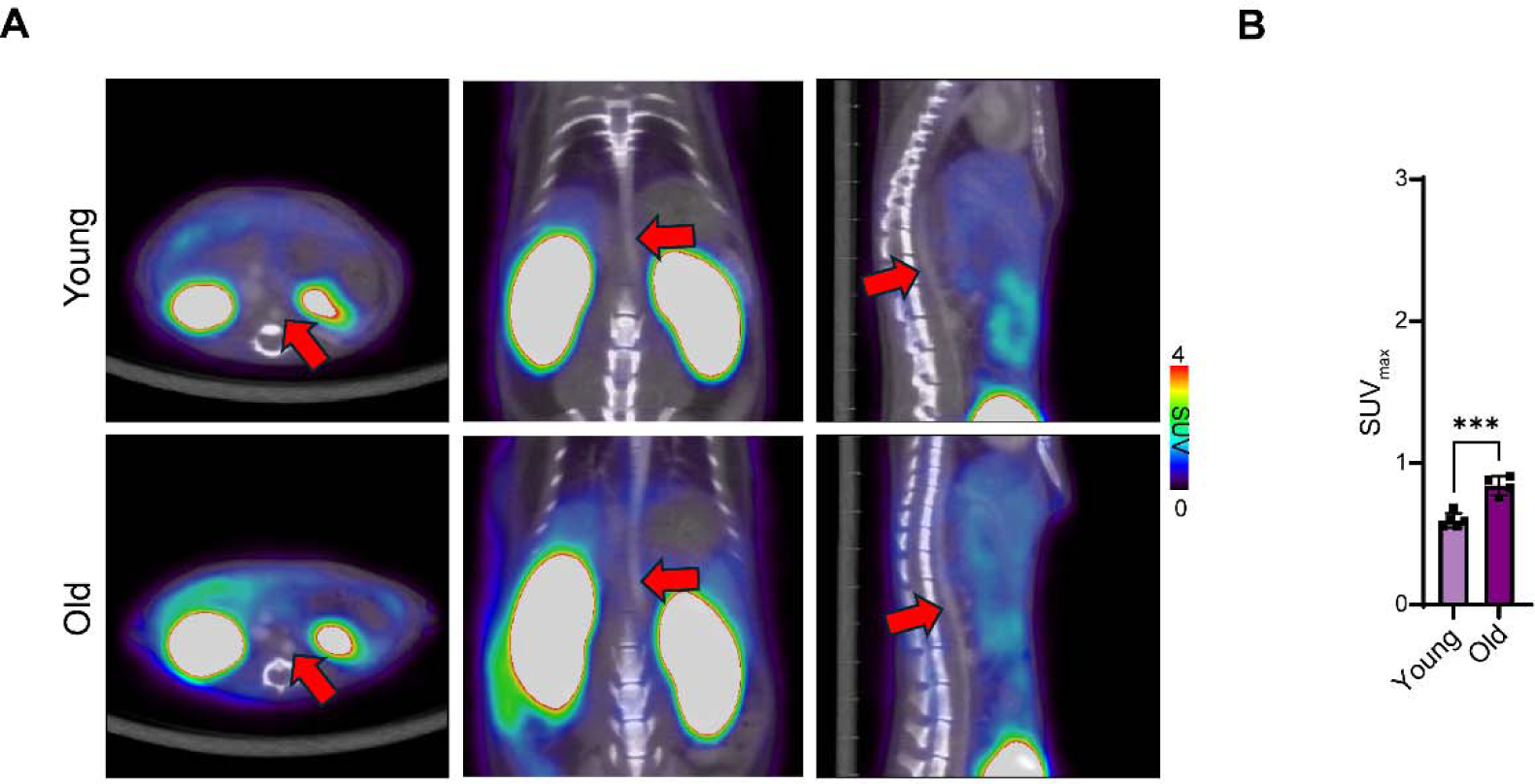
Age and ^64^Cu-RYM2 PET/CT in control *Apoe^−/−^* mice. **A.** Illustrative examples of fused ^64^Cu-RYM2 PET/contrast-enhanced CT images (axial, coronal, and sagittal) acquired in control young (8–10 weeks) and old (>51 weeks) *Apoe^−/−^* mice. Red arrows indicate the suprarenal abdominal aorta. **B.** Quantification of the ^64^Cu-RYM2 signal in the suprarenal abdominal aorta in young and old mice. ***: *P* < 0.001. SUV: Standardized uptake value.

### MMP activity and aortic dissection

The development of aortic aneurysm following Ang II infusion is preceded by focal medial dissections manifesting as intramural hematoma, which ultimately evolve into aneurysmal aortic remodeling (*18*). To investigate the time course of increased aorta MMP activity following Ang II infusion and whether this increase precedes or follows aortic remodeling, we evaluated MMP activity in fully mature, intermediate age (12- to 17-week-old) *Apoe^−/−^* mice before and after Ang II infusion, and verified key findings in younger and older animals. Tissue zymography showed a gradual increase in MMP activity with significantly higher levels on day 4 post-Ang II infusion compared to control animals (Figure 5). Notably, while there was seldom any evidence of dissection along the aorta before day 3, and especially day 4, showed early dissection as evidenced by the presence of intramural hematoma in the suprarenal abdominal aorta (Supplemental Figure 4). Together, these data suggest that increased MMP activity in the aorta precedes aortic dissection, an early stage in aneurysm development and rupture in this model.

**Figure 5.**
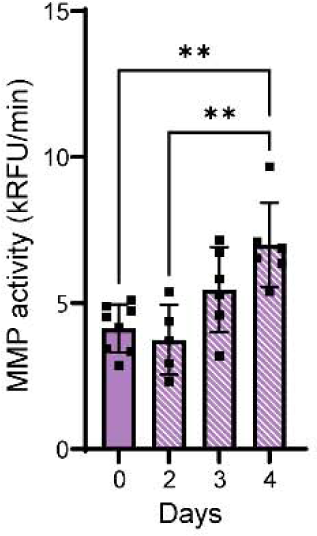
Aorta MMP activity in control (day 0) and angiotensin II-infused *Apoe^−/−^* mice (12-17 weeks) at different times after starting angiotensin II infusion. **: *P* < 0.01. RFU: Relative fluorescence units.

### Heterogeneity of MMP activity along the aorta

Given the SRA location of aortic dissection, we evaluated the pattern of MMP activity along the aorta before and early after Ang II infusion in 12- to 17-week-old *Apoe^−/−^* mice. To generate a detailed map of MMP activity along the aorta, we compared two molecular imaging approaches, fluorescence imaging using MMPSense 645 (a panMMP substrate that becomes fluorescent upon cleavage by MMPs) and autoradiography using ^99m^Tc-RYM1 (a ^99m^Tc-labled ^64^Cu-RYM2 homolog tracer, which similarly binds to the common MMP activation epitope) with tissue Zymography. Although all three techniques showed higher MMP activity in the ascending thoracic aorta (ATA), only the difference detected by tissue zymography reached statistical significance (MMP activity in ATA compared to DTA (*P* < 0.01), SRA (*P* < 0.01) or IRA (*P* < 0.01, Supplemental Figures 5 and Figure 6A). The MMP activity was also higher in ATA compared to other segments of the aorta in animals treated with Ang II for 3 days (MMP activity in ATA compared to DTA (*P* < 0.0001), SRA (*P* < 0.0001) or IRA (*P* < 0.0001, Figure 6B). However, the Ang II-induced increase in MMP activity was not uniform along the aorta, with the ATA and SRA, but not DTA and IRA, showing significant increases in MMP activity on day 3 post-Ang II infusion (*P* < 0.05 for ATA and SRA, Figure 6C). To investigate the effect of aging on the heterogeneity of MMP activity along the aorta, similar studies were performed in 8-10 week- and >51-week-old *Apoe^−/−^*mice. The higher ATA MMP activity was also detectable in 8–10-week-old animals. Additionally, aging was associated with a significant increase in MMP activity in ATA (*P* < 0.05, Supplemental Figure 6).

**Figure 6.**
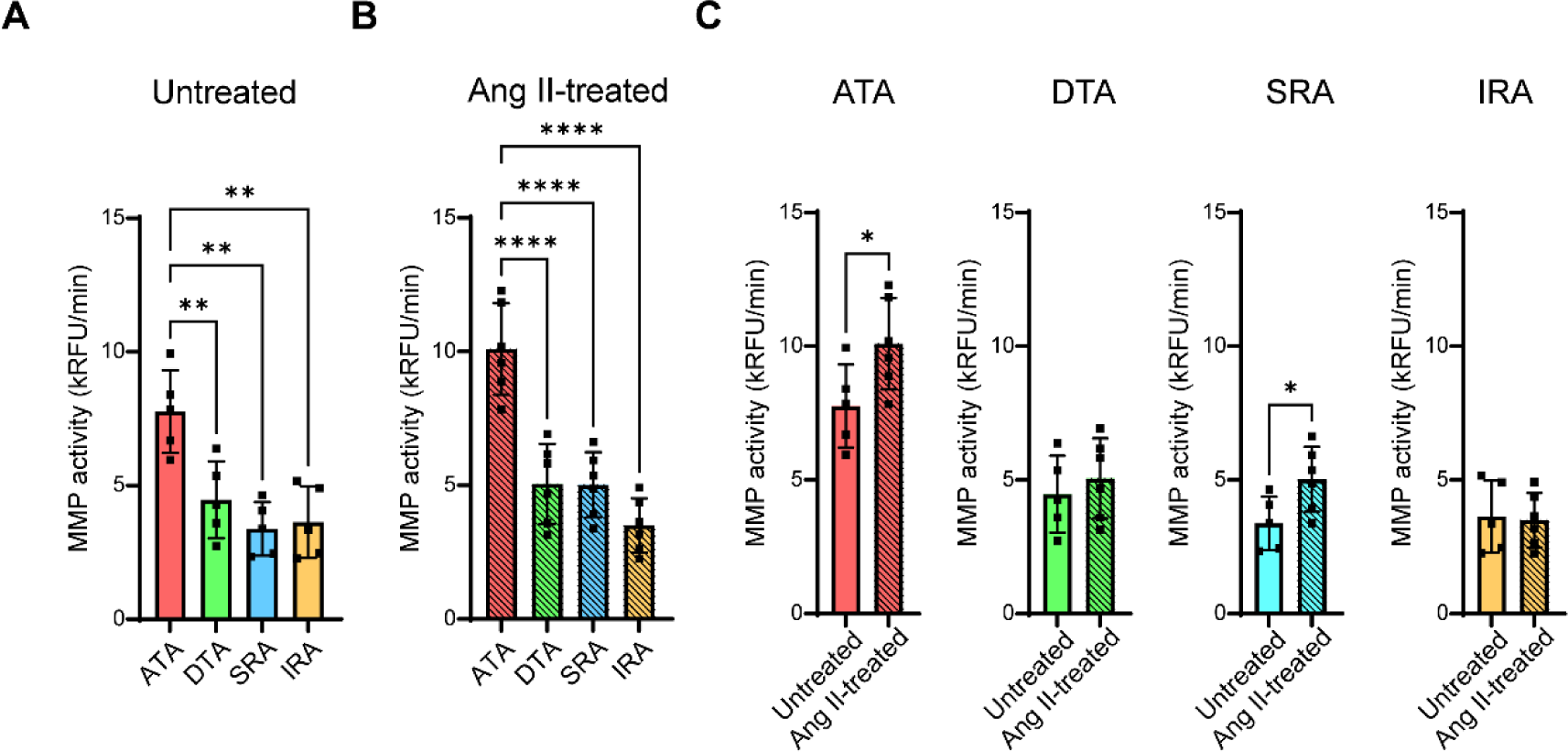
MMP activity in different segments of the aorta of *Apoe^−/−^* mice (12-17 weeks). **A, B.** MMP activity in different aorta segments from untreated *Apoe^−/−^* mice (**A**), and *Apoe^−/−^* mice treated with angiotensin II for 3 days (**B**). **C.** Comparison of MMP activity between untreated and angiotensin II-treated mice within each segment of the aorta. *:*P* < 0.05, **: *P* < 0.01, ****: *P* <0.0001. RFU: Relative fluorescence units, ATA: Ascending thoracic aorta, DTA: Descending thoracic aorta, SRA: suprarenal abdominal aorta, IRA: Infrarenal abdominal aorta.

### MMP expression along the aorta

Evaluation of selective members of the MMP family showed no significant differences in *Mmp2, 3, 9, and 13* gene expression in ATA compared to the other 3 segments of the aorta in 12- to 17-week-old *Apoe^−/−^*mice (except for ATA vs IRA for *Mmp2*). However, *Mmp12* expression was significantly higher in ATA than in the other three segments of the aorta (Supplemental Figures 7A-11A). On the other hand, the increase in ATA and SRA MMP activity at 3 days post Ang II infusion was associated with significantly higher *Mmp2* gene expression in ATA compared to DTA and IRA, and *Mmp13* expression in ATA compared to DTA and IRA, and SRA compared to IRA, while there was no significant change in *Mmp3*, *9*, and *12* expression across different aorta segments (Supplemental Figures 7B-11B). Highlighting the segmental variation in aortic response to Ang II infusion, while Ang II infusion significantly increased *Mmp2* and *13* expression in ATA and *Mmp13* in SRA, there was no increase in *Mmp2* and *13* (or any other MMP studied) gene expression in DTA and IRA (Supplemental Figures 7-11).

Next, we investigated the association between MMP activity and gene expression with aortic dissection/hematoma. Notably, MMPSense 645 imaging showed a focal increase in MMP activity in Ang II-induced SRA hematomas (Supplemental Figure 4). Evaluation of SRA MMP gene expression between segments with and without dissection (collected on days 3 and 4 of Ang II infusion) showed significantly higher expression of *Mmp3*, *9*, *12*, and *13* in dissected segments compared to the adjacent, apparently normal segments of the aorta (Figure 7).

**Figure 7.**
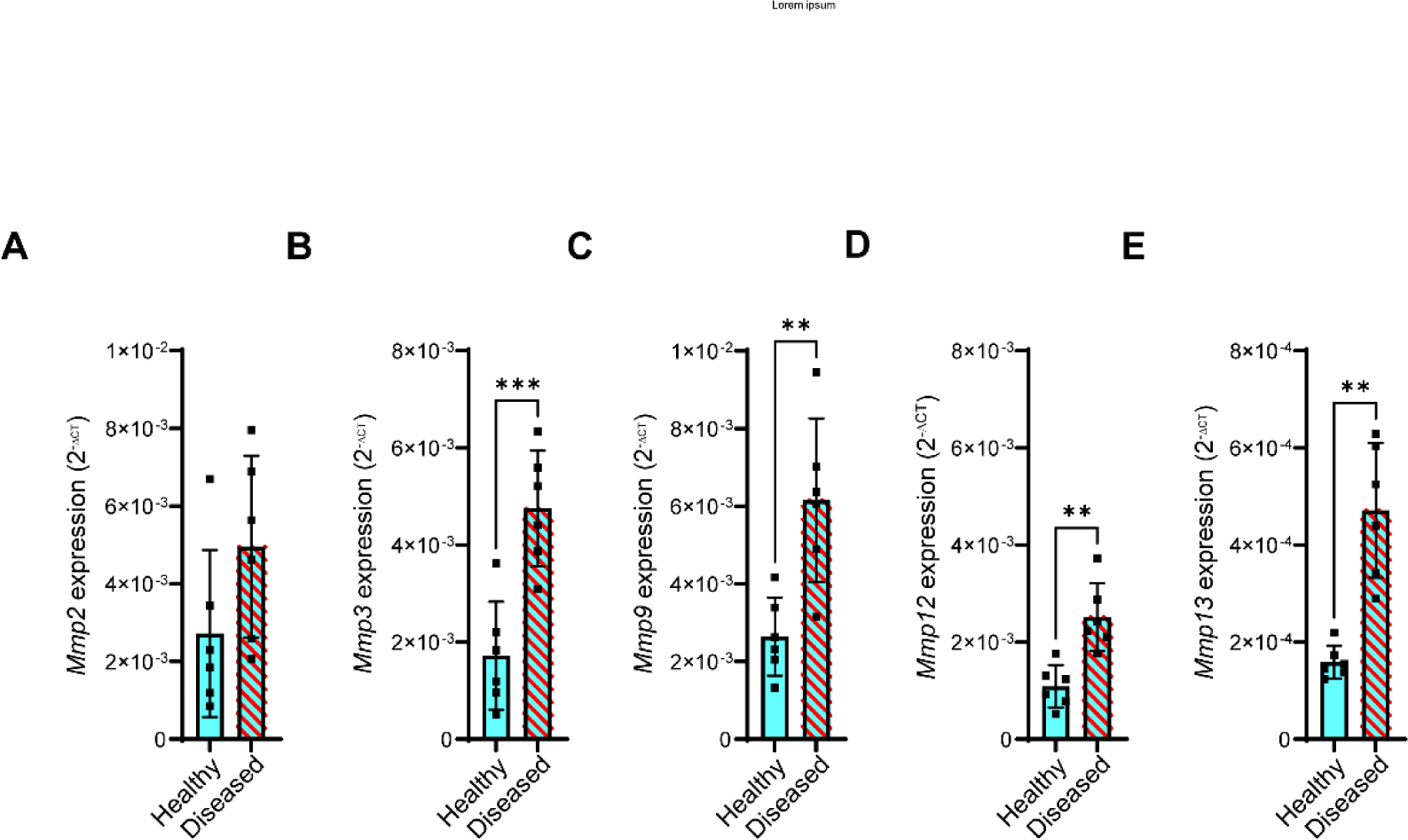
MMP expression in the suprarenal abdominal aorta. **A**-**E**) RT-qPCR analysis of *Mmp2* (**A**), *Mmp3* (**B**), *Mmp9* (**C**), *Mmp12* (**D**), and *Mmp13* (**E**) expression in apparently healthy (non-affected) and diseased (hematoma/dissected) portions of the suprarenal abdominal aorta of *Apoe^−/−^* mice (12-17 weeks) infused with angiotensin II for 3 or 4 days. Gene expression is reported as 2^-ΔCt^ relative to *Gapdh*. **: *P* < 0.01, ***: *P* < 0.001.

### Aging and the heterogeneity of MMP activity along the aorta in WT mice

To investigate whether a similar heterogeneous pattern of MMP activity exists in WT mice, we evaluated MMP activity by tissue zymography in 8-10 (young) and >51 (old) week-old WT mice. While there was no difference in MMP activity between different segments of the aorta in young mice, older WT mice showed a heterogeneous MMP activity pattern along the aorta, with significantly higher MMP activity in ATA compared to the other three segments of the aorta (*P* < 0.01, 0.05 and 0.01, respectively for DTA, SRA and IRA, Figure 8A and B). The comparison of MMP activity across different aortic segments between these two groups of animals showed a trend toward increased MMP activity with aging, which was statistically significant in ATA (*P* < 0.05, Figure 8C). A comparison of MMP activity between WT and *Apoe^−/−^* mice showed higher MMP activity in the ATA of *Apoe^−/−^* mice in both young and old animals. In young mice, the MMP activity was also significantly higher in the SRA of *Apoe^−/−^* mice compared to WT animals (Supplemental Figure 12).

**Figure 8.**
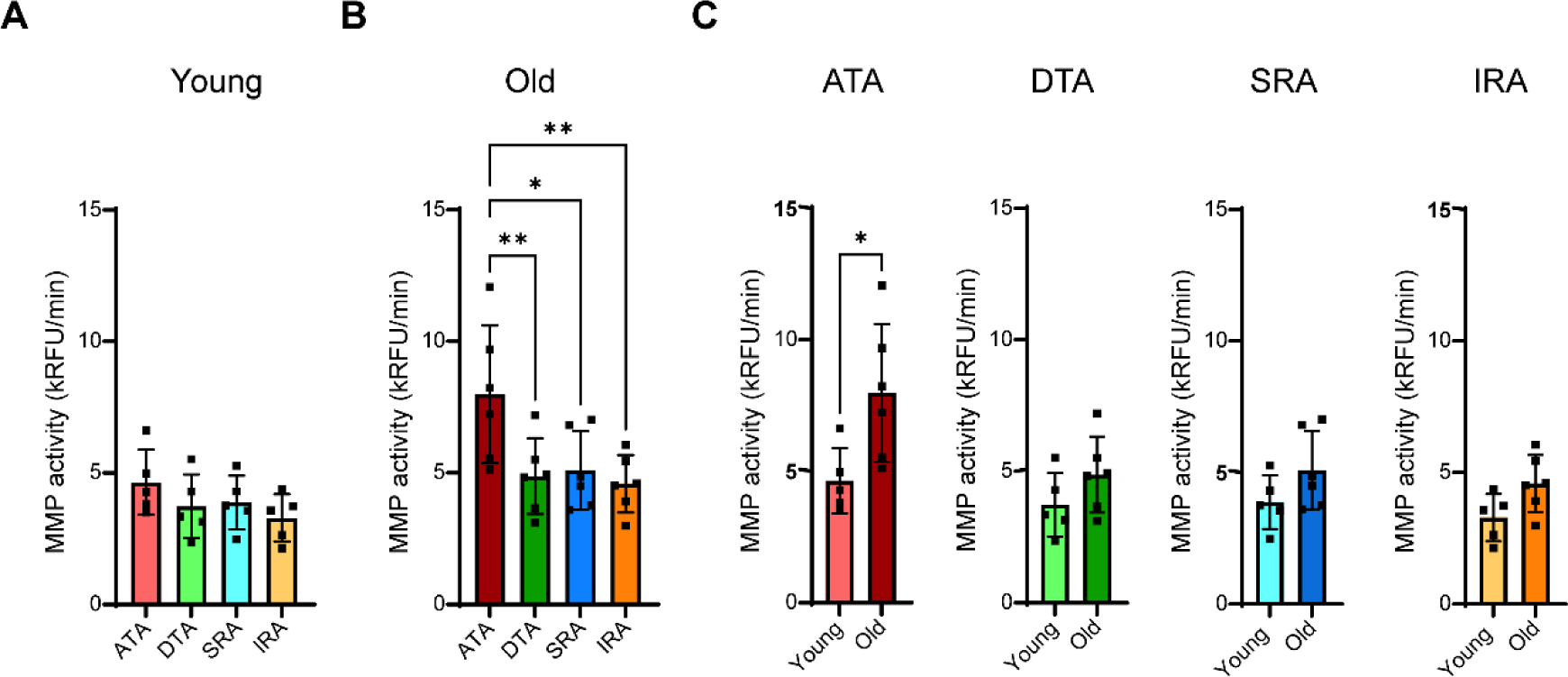
MMP activity in different segments of the aorta of wild-type mice. **A, B.** MMP activity in different aorta segments from young (8–10 weeks, **A**), and old (>51 weeks) mice (**B**). **C.** Comparison of MMP activity between young and old mice within each segment of the aorta. *: *P* < 0.05, **: *P* < 0.01. RFU: Relative fluorescence units, ATA: Ascending thoracic aorta, DTA: Descending thoracic aorta, SRA: suprarenal abdominal aorta, IRA: Infrarenal abdominal aorta.

## Discussion

This study offers new insight into the role of MMPs in aortic aneurysm. Aging is linked to decreased survival in mice infused with Ang II to induce aortic aneurysm, as well as increased aortic MMP activation detected by molecular imaging before and after Ang II infusion. The aortic MMP signal in early in vivo PET/CT images after Ang II infusion indicates survival prospects. Additionally, aging is associated with increased MMP activity and its heterogeneity along the aorta. Ang II infusion differentially promotes MMP activity along the aorta, and aortic dissection is associated with elevated MMP activity, with different MMP family members contributing to these changes before and after dissection.

Advanced age is a risk factor for the sporadic form of TAA and AAA (*2,19*). Age may also influence aneurysm evolution. While the human data on TAA in this regard are inconsistent (*2*), the incidence of rupture increases with age in AAA (*20*). Only a small number of preclinical studies have formally examined the impact of age on AAA formation and outcomes in controlled settings (*11*). A study comparing AAA development in Ang II-infused young (10-week-old) and old (24-week-old) *Apoe^−/−^* mice reported a greater AAA-to-normal aorta diameter ratio in the old group (*21*). Another study reported a higher incidence of AAA and lower survival in Ang II-infused C57Bl/6 mice aged 18 to 20 months than those aged 2-3 months (*22*). This was associated with increased aortic MMP-2 expression in older animals, which was linked to reduced sirtuin 1 (a histone deacetylase) expression and activity, and increased vascular cell senescence and vascular inflammation (*22*). Our study confirms and expands the scope of these findings, demonstrating a significant reduction in survival in old compared to young animals, which is associated with enhanced MMP activation detected by molecular imaging.

MMPs are upregulated in human and murine aortic aneurysms, and several studies have indicated higher MMP expression in aortic aneurysms compared to the normal aorta (*12,13*). Molecular imaging can provide quantitative information regarding MMP activation along the aorta, enabling longitudinal studies in the same animal. We leveraged MMP-targeted molecular imaging to investigate the effect of animal age on aortic MMP activation in control and Ang II-infused mice and its relation to aneurysm outcome. ^64^Cu-RYM2 (a tracer that binds to the common MMP activation epitope) (*13*) PET/CT imaging showed higher aorta tracer uptake in surviving old mice than in young animals after 1 week of Ang II infusion. A similar difference in MMP tracer uptake between old and young mice was detected in control animals, with Ang II infusion increasing tracer uptake in both groups. Notably, SRA tracer uptake was significantly higher in animals that did not survive 28 days than in those that did. This indicates that aortic MMP activation can serve as a marker and potentially contributes to aneurysm development and rupture.

A few studies have pointed to differences in MMP expression between different regions of the aorta and in surgical specimens collected after aortic dissection or aneurysm rupture (*12,23*). However, no systematic evaluation of MMP activity and expression along the aorta has been reported. The aorta is a complex organ with significant regional differences, including its structural and biomechanical properties (*23*). The decrease in the aorta size along its path is associated with key differences in the number of lamellar units separated by elastic laminae between the thoracic and abdominal aorta, which respectively contain 55-60 and 28-32 lamellar units in humans (*23*). These differences lead to changes in aortic stiffness, which gradually increases along the aorta (*24,25*). The transition from turbulent to laminar flow, the change in blood flow direction in the aortic arch, and the presence of side branches along the descending aorta also contribute to the biomechanical heterogeneity of the aorta. Smooth muscle cells contribute to the integrity of the aortic wall, and their loss is a key feature in the pathogenesis of AAA. There are significant differences in the embryologic origins of SMC along the aorta, with those in the proximal thoracic aorta (including the aortic arch) deriving from the neural crest and the secondary heart field. In contrast, the SMC in the rest of the aorta are mesodermal derivatives (*26,27*). Combined with the aorta’s structural and biomechanical heterogeneity, these regional differences in SMC contribute to the varying susceptibility of the aorta to aneurysm and the differences in aneurysm pathobiology across the aorta (*23*). As such, AAA is often located in the infrarenal aorta in humans and is frequently associated with atherosclerosis, whereas single-gene mutations in extracellular matrix proteins, SMC contractility proteins, and mediators of TGFβ signaling play a more prominent role in TAA (*28*).

The analysis of MMP activity and expression along the mouse aorta shows a similar heterogeneity, which becomes more pronounced with aging, Ang II infusion, and dissection. As such, while there is no difference in MMP activity across different segments of the aorta in young wild-type mice, aging is associated with significantly higher MMP activity in the ATA. The higher MMP activity in the ATA is also observed in *Apoe^−/−^* mice, regardless of their age. *Apoe^−/−^*mice are prone to developing atherosclerosis, and we cannot exclude the possibility that the higher ATA MMP activity observed in older *Apoe^−/−^*mice is in part related to atherosclerosis in this atheroprone segment of the aorta. However, the higher ATA MMP activity in old wild-type mice links aging to increased ATA MMP activity in the absence of atherosclerosis. The higher MMP activity in *Apoe^−/−^* mice compared to wild-type animals may also contribute to the higher propensity to aneurysm development in these animals (*11*). Ang II infusion in *Apoe^−/−^*mice leads to an increase in MMP activity within 3-4 days that is limited to ATA and SRA, segments that have a higher predilection for aneurysm formation, dissection, and rupture (*11,29,30*). The higher predilection of SRA relative to ATA for developing dissecting aneurysms may be linked to an early mural tear near the ostium of small suprarenal side branches and local destruction of the medial architecture with or without intramural hematoma near the ostium of small abdominal side branches, as detected by differential phase contrast X-ray tomographic microscopy and histology (*31*). Among the MMPs evaluated, this increased activity is associated with upregulation of *Mmp2* and *Mmp13* in the ATA and of *Mmp2* in the SRA. Moreover, the earliest evidence of aortic aneurysm and dissection, i.e., intramural hematoma, is detected in SRA on day 3-4 of Ang II infusion, suggesting that MMP activation in this segment may set the stage for subsequent events. Notably, MMP activity is significantly higher in the dissected part of the SRA than in its non-affected segment, suggesting that dissection promotes MMP activity. This increased MMP activity in dissected SRA is associated with the upregulation of several members of the MMP family, including *Mmp3*, *9*, *12*, and *13*, highlighting a distinct MMP signature of dissection in aortic aneurysms.

## Limitations

We cannot rule out the possibility that differences in tracer clearance have contributed to the difference in the MMP signal between young and old animals on PET/CT images. However, a higher MMP activity in old animals was also observed with an independent technique, tissue zymography. Our studies of selected members of the MMP family revealed notable differences in MMP expression along the aorta. Different MMPs play distinct roles in vascular biology, and other MMPs (as well as tissue inhibitors of MMPs) may contribute to the heterogeneity of the aorta and to the development, dissection, and rupture of aortic aneurysms. MMP activity depends not only on MMP expression but also on MMP activation (and the presence of MMP inhibitors). While there was no significant difference in the expression of some MMP family members along the aorta, we cannot exclude changes in their activation (and biological activity) without a corresponding change in MMP expression. Interestingly, it has been reported that aging is associated with increased MMP-2 activity in the thoracic aorta (most likely the ascending thoracic aorta) in humans (*15*). In the absence of radiotracers and other agents highly selective for specific MMPs, we could only evaluate overall MMP activation in our longitudinal studies of MMP activation and aneurysm outcomes.

## Conclusions

Aging is associated with increased MMP activity and heterogeneity along the aorta, which may underlie the worse aneurysm survival in older animals. Highlighting the role of MMPs in aneurysm pathobiology, MMP activation detected by molecular imaging can inform the aneurysm survival prospects. Taken together, these data suggest that inhibitors and tracers targeting specific members of the MMP family may help prevent and track selective aspects of aneurysm growth, dissection, and rupture.

## KEY POINTS

QUESTION: How does aging affect aortic MMP expression and activity, as well as aneurysm development and survival?

PERTINENT FINDINGS: PET/CT using ^64^Cu-RYM2, an MMP-targeted radiotracer, showed higher tracer uptake in the aorta of old mice compared to young animals before and 7 days after angiotensin II infusion. It also identified animals with lower survival prospects. Tissue zymography revealed a heterogeneous pattern of MMP expression and activity along the aorta, which increased with aging and angiotensin II infusion.

IMPLICATIONS FOR PATIENT CARE: Inhibitors and tracers targeting specific members of the MMP family may help prevent and track selective aspects of aortic aneurysm growth, dissection, and rupture.

