## Supplemental Data for "Aging, matrix metalloproteinase imaging, and survival prospects in aortic aneurysm"

### Supplemental Material

### Table of Contents


| | |
|---|---|
| Detailed Materials and Methods | 3 |
| Supplemental Figure 1 | 7 |
| Supplemental Figure 2 | 8 |
| Supplemental Figure 3 | 9 |
| Supplemental Figure 4 | 10 |
| Supplemental Figure 5 | 11 |
| Supplemental Figure 6 | 12 |
| Supplemental Figure 7 | 13 |
| Supplemental Figure 8 | 14 |
| Supplemental Figure9 | 15 |
| Supplemental Figure10 | 16 |
| Supplemental Figure11 | 17 |
| Supplemental Figure12 | 18 |

#### Supplemental Materials and Methods

##### RYM2 labeling

Radiolabeling of RYM2 with $^{64}Cu$ was performed following previously reported procedures (*1*). Briefly, cyclotron-produced $^{64}CuCl_2$ (Washington University School of Medicine Mallinckrodt Institute of Radiology Cyclotron Facility) was converted to $^{64}Cu(OAc)_2$ by adding 50 μL of 0.5M NaOAc (pH3.5). RYM2 (5 nmol in 5 μL DMSO) was mixed with 60 μL of 0.1M NaOAc (pH 5.5) and added to the radioisotope and heated for 30 min at 45 °C. The radiolabeled solution was cooled to room temperature and subjected to quality control using radio high-performance liquid chromatography (HPLC). HPLC separation was carried out on a Phenomenex C18 reverse-phase column (Luna 5 µm C18(2), 100 Å, LC Column 250 x 4.6 mm). Gradient elution was performed using water with 0.1% trifluoacetic acid (TFA) as mobile phase A (A) and acetonitrile with 0.1% trifluoacetic acid (TFA) as mobile phase B (B). Flow rate was set at 1 mL/min, 5-95% B gradient in 10 min, a 1-min isocratic phase, which was linearly brought back to the initial gradient by the end of the run. Chromatograms were monitored by ultraviolet detection at 254 and 214 nm and by in-line radioactivity detection for $^{64}Cu$-labeled compounds. The specific activity of $^{64}Cu$-RYM2 was 78.2 ± 8.4 GBq/µmol.

##### PET/CT imaging

$^{64}Cu$-RYM2 PET/CT was performed as described (*1*). Briefly, under inhaled isoflurane anesthesia, the animals were injected intravenously with 11.73 ± 3.50 MBq of $^{64}Cu$-RYM2 and 30 µL + (body weight in grams) µL of Exitron nano 12000 (Viscover Imaging). A CT acquisition was then carried out, followed by a 10-minute PET acquisition on a small-animal dedicated PET/CT scanner (Inveon PET/CT, Siemens Medical Solutions), beginning 50 minutes after tracer injection. CT images for attenuation correction were reconstructed at a down-sampled resolution (0.815×0.815×0.796 mm) using the Feldkamp cone-beam algorithm. CT images for co-registration with PET images were reconstructed using a 3D ordered-subset expectation maximization (OSEM3D) algorithm with an isotropic voxel size of 0.111 mm.

Emission data were reconstructed using the manufacturer’s software (Inveon Workstation, version 2.0) with the OSEM3D algorithm (2 iterations, 16 subsets), followed by a maximum a posteriori probability reconstruction (MAP; 25 iterations), producing images with a voxel size of 0.388×0.388×0.796 mm and incorporating decay, attenuation, scatter, normalization, and randoms correction. PET images were normalized to the injected dose and body weight and displayed on a standardized uptake value (SUV) scale. One Ang II-infused animal from the "old" group died during imaging, and its images were not analyzed.

**PET/CT image analysis**

Quantitative images analysis was performed using 3D Slicer software (version 5.2.2; https://www.slicer.org/) (*2*), and AMIDE software (version 1.0.6; https://amide.sourceforge.net/) was used to display images. Contrast-enhanced CT images were used to localize the aorta and draw volumes-of-interest (VOIs) over the aorta as follows: the descending thoracic aorta was from the aortic arch to the diaphragm, the suprarenal aorta was from the diaphragm to the renal arteries, and the infrarenal aorta was from the renal to the iliac arteries. These VOIs were then applied to the PET volume, ensuring they did not overlap with the spillover signal from the kidneys and adjacent uptake from the lumen area when present is included. Inter-observer variability was assessed using independent analyses of a subset of imaging data by two different observers (AA, NG). The intra-class correlation coefficient for the measurement was 0.87, confirming the reproducibility of the image analysis. A previously acquired PET/CT dataset of young *Apoe*$^{-/-}$ mice (n = 5) (*1*) was reanalyzed for use in the current study. Image analysis of this dataset was independently repeated using the same workflow, and the reanalysis measurements closely matched the original values (intra-class correlation coefficient = 0.98).

**Near infrared fluorescence imaging**

MMPSense 645 FAST (Revvity), an activatable pan-MMP probe that becomes fluorescent following cleavage by MMPs, was prepared according to the manufacturer's instructions and administered to animals intravenously at a dose of 160 nmol/kg. Twenty-four hours later, the animals were anesthetized. A solution of cold PBS was infused into the left ventricle, and the right atrium was incised to serve as outflow. The aorta was quickly cleaned of surrounding tissue under a stereomicroscope (MZ9.5, Leica), removed, and fixed with 4% PFA. The aorta was then opened longitudinally, flat-mounted, and imaged as a whole-mount preparation on a DMi8 widefield fluorescence microscope (Leica) using a compatible filter set. Autofluorescence images provided anatomical reference. ImageJ (NIH) was used to quantify the mean MMPSense 645 fluorescence intensity across different regions of interest (ROIs) defined in the autofluorescence image.

**Autoradiography**

Autoradiography was performed as previously described (*3*). Mice were injected intravenously with $^{99m}$Tc-RYM1 (26.8 ± 2.3 MBq). Two hours later, the animals were anesthetized, and the aorta was dissected from surrounding tissues, placed on a phosphor imaging plate (MultiSensitive Phosphor Screen; PerkinElmer) with standards of known activity, and scanned with a high-resolution phosphor imager (Typhoon Trio; GE Healthcare Life Sciences). Digital images were analyzed in ImageJ (NIH) by drawing ROIs over predefined anatomical segments of the aorta. Tracer uptake was expressed as decay-corrected percent injected dose per square centimeter (%ID/cm$^2$).

**Morphometry**

5 µm-thick serial cross-sections of the abdominal aorta were used for morphometric analysis. The circumference of the external elastic lamina in each section was measured, and the maximal diameter was calculated, assuming a circular geometry.  A maximal diameter of > 1.5 mm defined the presence of an aortic aneurysm.

**H&E staining**

OCT-embedded aorta segments were sectioned at 5 µm thickness and mounted onto glass slides. OCT was removed by washing sections in PBS, and the tissue was fixed in 10% neutral buffered formalin for 15 minutes. Slides were stained with hematoxylin (Abcam) for 15 seconds, rinsed in two changes of distilled water, and incubated with a bluing reagent (Abcam) for 10 seconds. They were then briefly dipped in absolute alcohol, counterstained with modified alcoholic eosin Y (Abcam) for 3 minutes, rinsed in absolute alcohol, dehydrated through three changes of absolute alcohol, and mounted. Brightfield images were acquired with a light microscope (Bx63; Olympus).

**Tissue zymography**

Animals were anesthetized and perfused with cold PBS, as above. The aorta was then cleaned quickly and excised from surrounding tissues, cut into predefined anatomical segments, and embedded in OCT. The ascending thoracic aorta (ATA), which included the aortic arch, was the segment proximal to the left subclavian artery. The descending thoracic aorta (DTA) was the segment immediately after the left subclavian artery until the diaphragm. The suprarenal abdominal aorta (SRA) was the segment between the diaphragm and the left renal artery, and the infrarenal abdominal aorta (IRA) was the segment distal to renal arteries up to iliac bifurcation. Aortas exhibiting any evidence of hematomas or dissection (1 in the whole-aorta Ang II-induced MMP activity time-course study and 1 in the regional 3-day Ang II-induced MMP heterogeneity study) were excluded from analysis. The tissue MMP activity was detected and quantified in 1 µg of protein lysate as previously described (*3*) using a commercial fluorometric MMP activity assay, the SensoLyte 520 Generic MMP Activity Kit from AnaSpec, according to the manufacturer's instructions. MMP activity was measured at an excitation wavelength of 490 nm and an emission wavelength of 520 nm using Agilent BioTek Synergy H1 or Applied Biosystems 7500 fluorescence detection systems. Cross-instrument normalization was performed using a correction factor was derived from biologically matched reference samples.

**Gene expression analysis**

Total RNA was isolated from aortic tissues, and cDNA was synthesized using the RNeasy Plus Mini Kit and the QuantiTect Reverse Transcription Kit (Qiagen) according to the manufacturer's instructions. Real-time quantitative PCR was performed on a 7500 Applied Biosystems Real-Time PCR System (Thermo Fisher Scientific) using TaqMan Gene Expression Assays (Thermo Fisher Scientific). The following probes were used in this study: *Mmp2* (Mm00439498_m1), *Mmp3* (Mm00440295_m1), *Mmp9* (Mm00442991_m1), *Mmp12* (Mm00500554_m1), *Mmp13* (Mm00439491_m1), and glyceraldehyde-3-phosphate dehydrogenase (*Gapdh*, Mm99999915_g1). Gene expressions were reported as $2^{-\Delta Ct}$ relative to *Gapdh*).

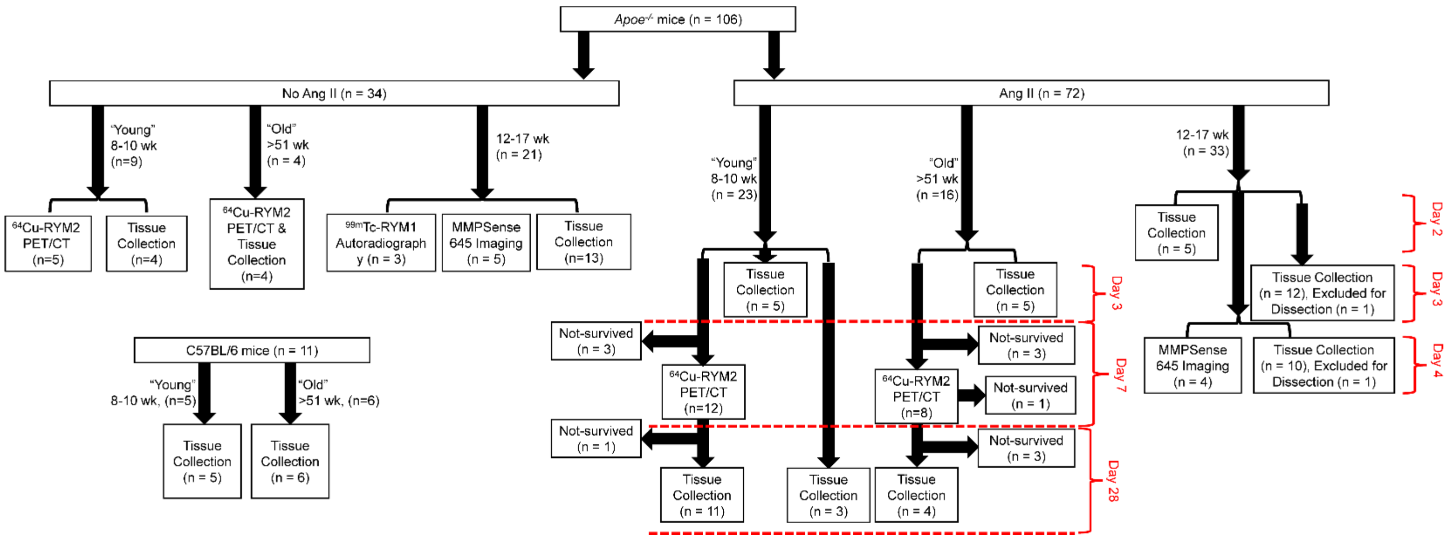


**Supplemental Figure 1.** Flow chart of animals used in the study and their endpoints. Animal survival was tracked in young (8-10 weeks) and old (>51 weeks) *Apoe*$^{-/-}$ mice infused with angiotensin II (Ang II) for up to 28 days. The animals underwent in vivo $^{64}$Cu-RYM2 PET/CT imaging after 7 days to evaluate aortic MMP activation in relation to survival to 28 days. The temporal and spatial patterns of MMP expression and activity and their relation to aortic dissection, an early step in aneurysm development in this model, was investigated by tissue zymography, ex vivo molecular imaging, and gene expression analysis in an intermediate age, fully adult (12-17 weeks) group of *Apoe*$^{-/-}$ mice infused with Ang II for up to 4 days and the effect of aging on these findings was corroborated in aortas collected from young and old *Apoe*$^{-/-}$ mice after 3 days of Ang II infusion. Age-matched non-Ang II-infused *Apoe*$^{-/-}$ mice and young (8-10 weeks) and old (>51 weeks) C57BL/6 mice were used as controls in different experiments.

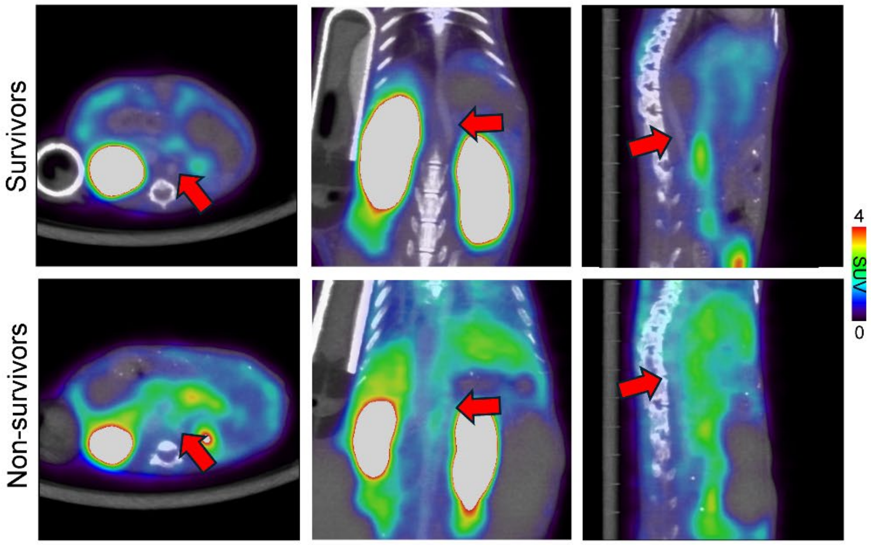


**Supplemental Figure 2.** Illustrative examples of fused $^{64}$Cu-RYM2 PET/contrast-enhanced CT images (axial, coronal, and sagittal) acquired 7 days after starting angiotensin II infusion in old (>51 weeks) mice that survived to day 28 (survivors) and from mice that died before day 28 (non-survivors). Red arrows indicate the suprarenal abdominal aorta.

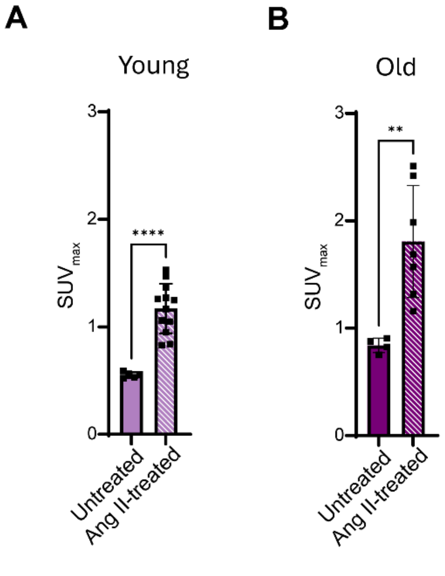


**Supplemental Figure 3.** Quantification of the suprarenal abdominal aorta $^{64}$Cu-RYM2 signal on PET/CT images acquired in untreated, and 7 days angiotensin II-treated young (8-10 weeks, **A**) and old (>51 weeks, **B**) *Apoe*$^{-/-}$ mice. **: $P < 0.01$, ****: $P < 0.0001$. SUV: Standardized uptake value.

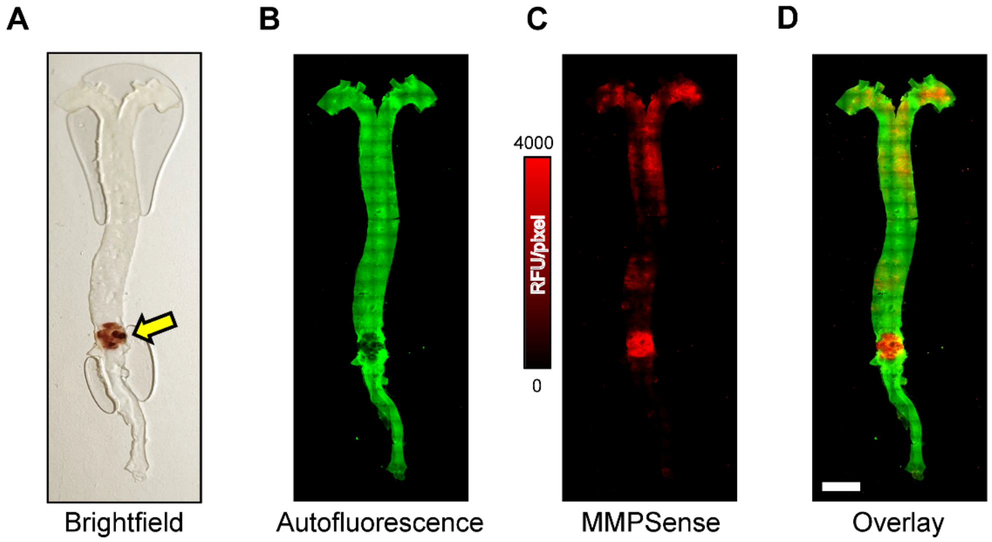


**Supplemental Figure 4.** Illustrative brightfield (**A**), autofluorescence (**B**), MMPSense 645 fluorescence (**C**), and fused MMPSense 645 fluorescence/autofluorescence (**D**) whole-mount images of the aorta of an angiotensin II-infused (for 4 days) *Apoe*$^{-/-}$ mouse (12-17 weeks) with intramural hematoma. Yellow arrow indicates the intramural hematoma. Scale bar = 2.5 mm.

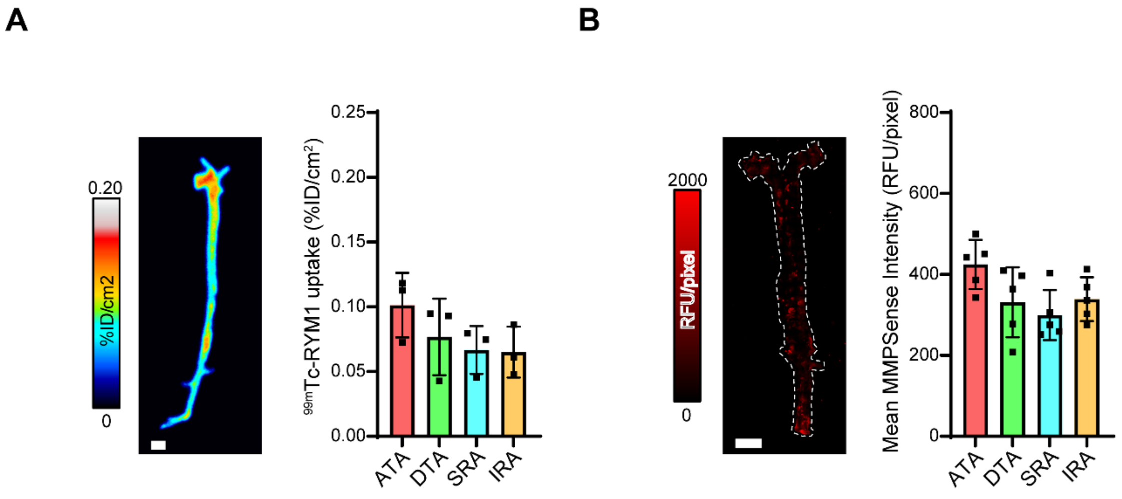


**Supplemental Figure 5.** Illustrative examples of $^{99m}$Tc-RYM1 autoradiography (**A**) and MMPSense 645 fluorescence imaging (**B**) and their quantification in different segments of the aorta of 12-17 weeks *Apoe*$^{-/-}$ mouse. Scale bar = 2.5 mm. ID: injected dose, RFU: Relative fluorescence units, ATA: Ascending thoracic aorta, DTA: Descending thoracic aorta, SRA: suprarenal abdominal aorta, IRA: Infrarenal abdominal aorta.

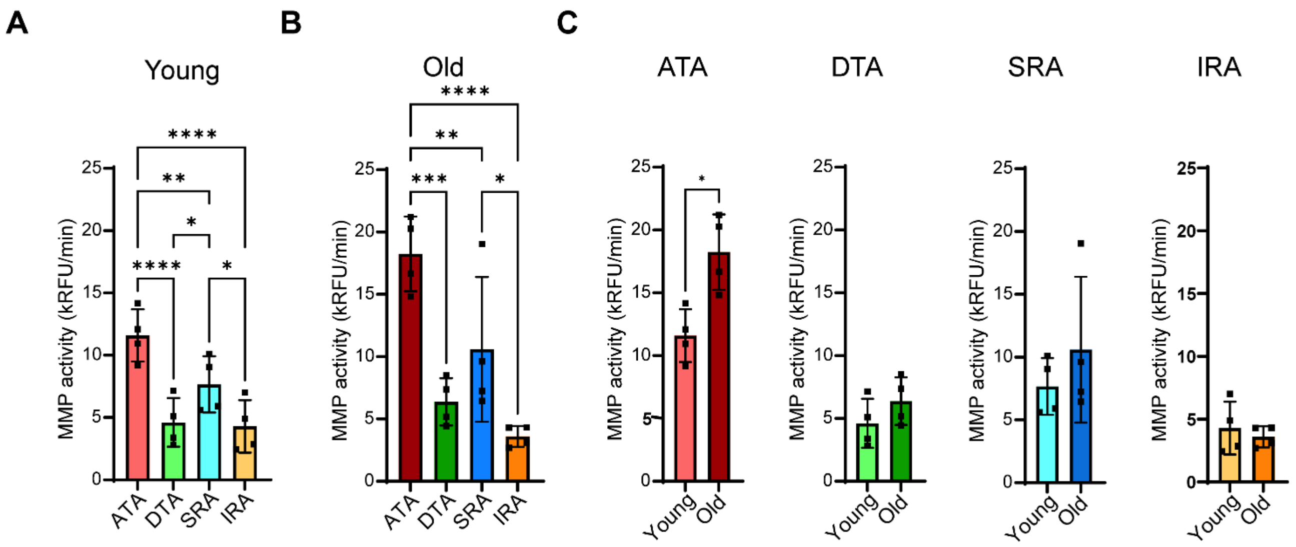


**Supplemental Figure 6.** MMP activity in different segments of the aorta of *Apoe*$^{-/-}$ mice. **A, B.** MMP activity in different aorta segments from young (8–10 weeks, **A**), and old mice (>51 weeks) mice (**B**). **C.** Comparison of MMP activity between young and old mice within each segment of the aorta. *: $P < 0.05$, **: $P < 0.01$, ***: $P < 0.001$, ****: $P < 0.0001$. RFU: Relative fluorescence units, ATA: Ascending thoracic aorta, DTA: Descending thoracic aorta, SRA: suprarenal abdominal aorta, IRA: Infrarenal abdominal aorta.

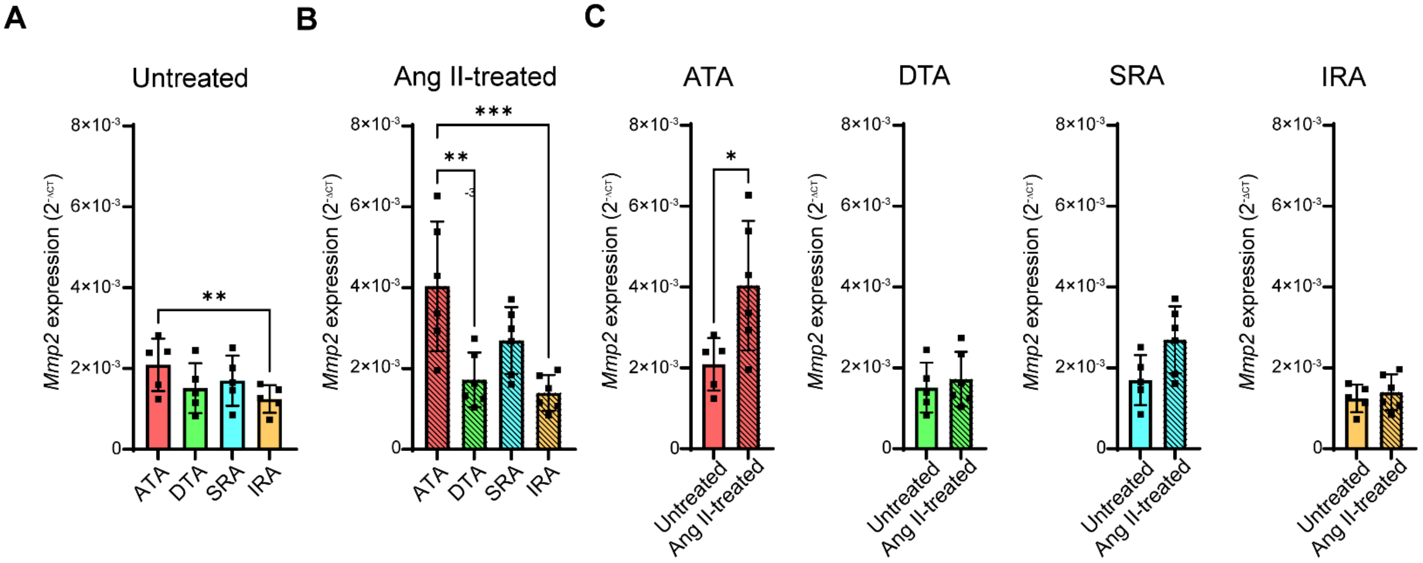


**Supplemental Figure 7.** *Mmp2* expression in different segments of the aorta of *Apoe*$^{-/-}$ mice. (12-17 weeks). **A, B.** *Gapdh*-normalized *Mmp2* expression in different aorta segments from untreated *Apoe*$^{-/-}$ mice (**A**), and *Apoe*$^{-/-}$ mice treated with angiotensin II for 3 days (**B**). **C.** Comparison of *Mmp2* expression between untreated and angiotensin II-treated mice within each segment of the aorta. *:$P$ < 0.05, **: $P$ < 0.01, ***: $P$ <0.001. RFU: Relative fluorescence units, ATA: Ascending thoracic aorta, DTA: Descending thoracic aorta, SRA: suprarenal abdominal aorta, IRA: Infrarenal abdominal aorta.

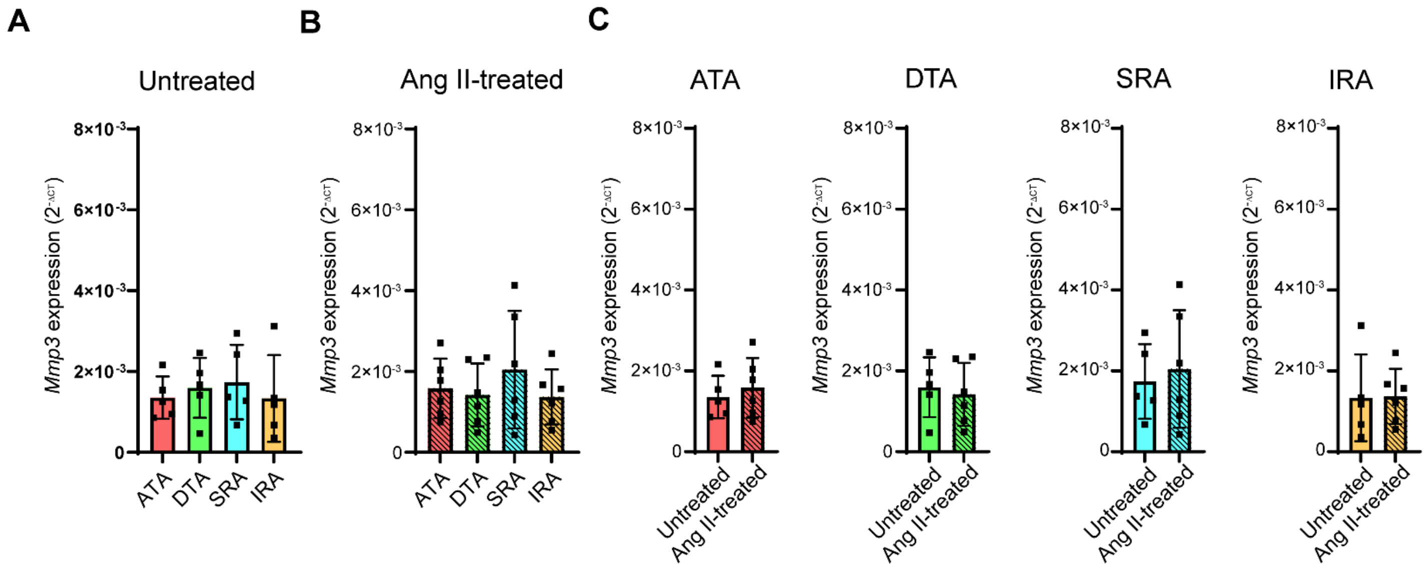


**Supplemental Figure 8.** *Mmp3* expression in different segments of the aorta of *Apoe*$^{-/-}$ mice. (12-17 weeks). **A, B.** *Gapdh*-normalized *Mmp3* expression in different aorta segments from untreated *Apoe*$^{-/-}$ mice (**A**), and *Apoe*$^{-/-}$ mice treated with angiotensin II for 3 days (**B**). **C.** Comparison of *Mmp3* expression between untreated and angiotensin II-treated mice within each segment of the aorta. RFU: Relative fluorescence units, ATA: Ascending thoracic aorta, DTA: Descending thoracic aorta, SRA: suprarenal abdominal aorta, IRA: Infrarenal abdominal aorta.

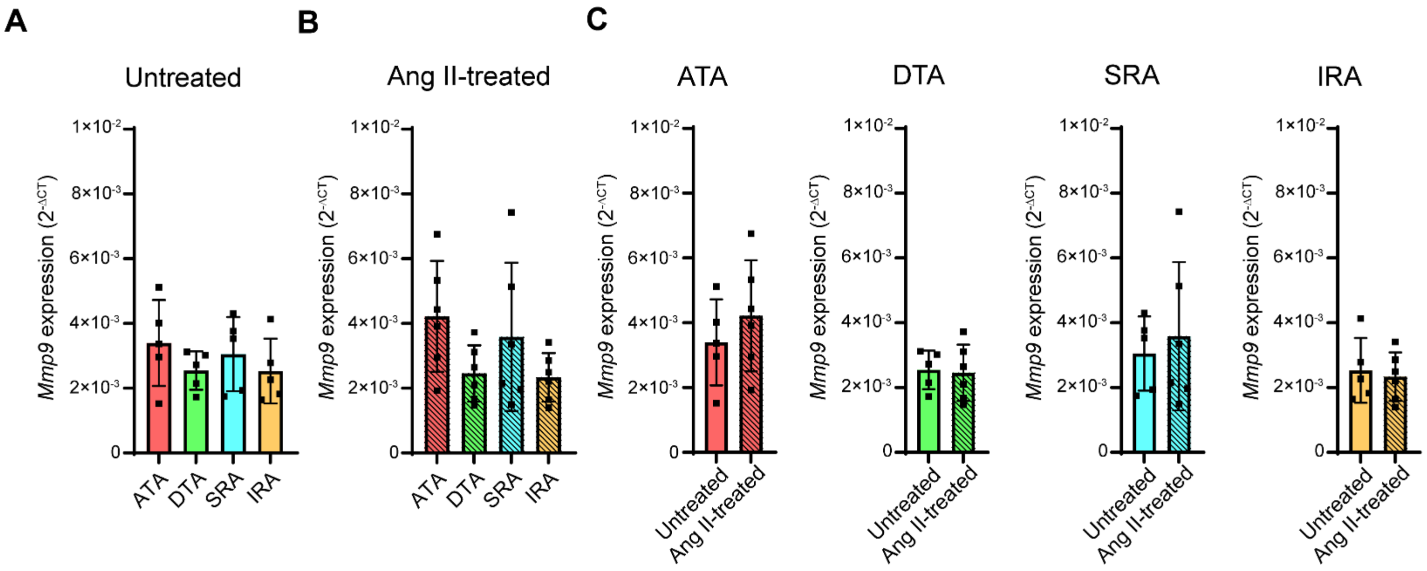


**Supplemental Figure 9.** *Mmp9* expression in different segments of the aorta of *Apoe*$^{-/-}$ mice. (12-17 weeks). **A, B.** *Gapdh*-normalized *Mmp9* expression in different aorta segments from untreated *Apoe*$^{-/-}$ mice (**A**), and *Apoe*$^{-/-}$ mice treated with angiotensin II for 3 days (**B**). **C.** Comparison of *Mmp9* expression between untreated and angiotensin II-treated mice within each segment of the aorta. RFU: Relative fluorescence units, ATA: Ascending thoracic aorta, DTA: Descending thoracic aorta, SRA: suprarenal abdominal aorta, IRA: Infrarenal abdominal aorta.

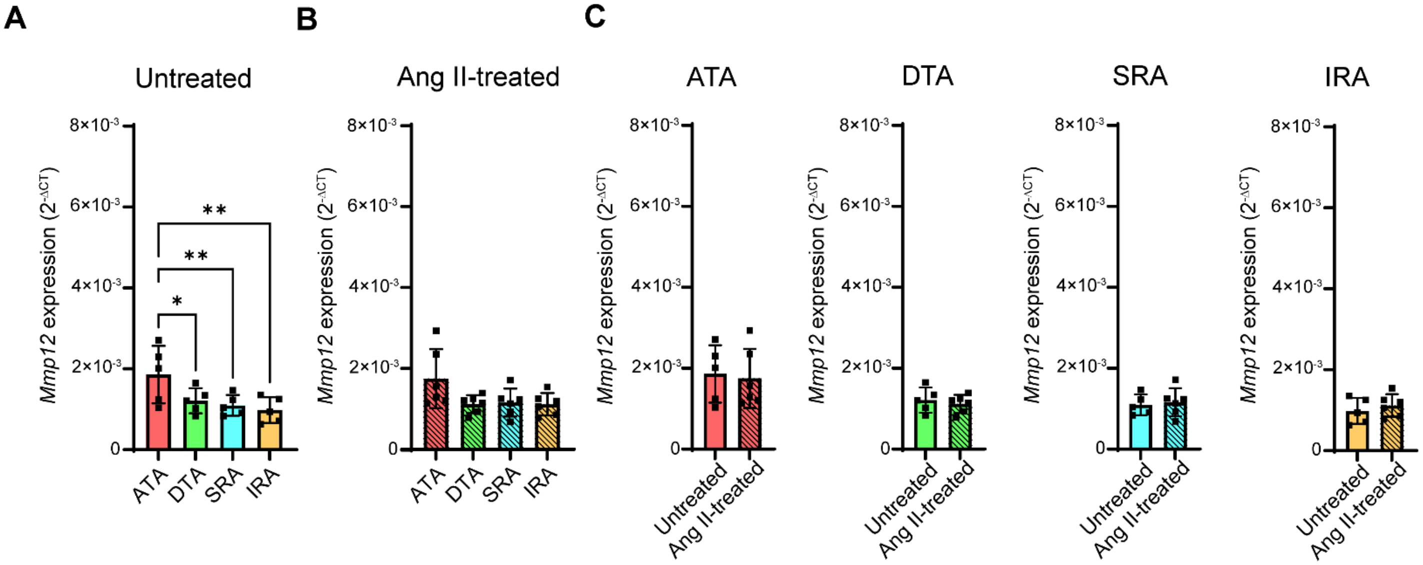


**Supplemental Figure 10.** *Mmp12* expression in different segments of the aorta of *Apoe*$^{-/-}$ mice. (12-17 weeks). **A, B.** *Gapdh*-normalized *Mmp12* expression in different aorta segments from untreated *Apoe*$^{-/-}$ mice (**A**), and *Apoe*$^{-/-}$ mice treated with angiotensin II for 3 days (**B**). **C.** Comparison of *Mmp12* expression between untreated and angiotensin II-treated mice within each segment of the aorta. *: $P < 0.05$, **: $P < 0.01$. RFU: Relative fluorescence units, ATA: Ascending thoracic aorta, DTA: Descending thoracic aorta, SRA: suprarenal abdominal aorta, IRA: Infrarenal abdominal aorta.

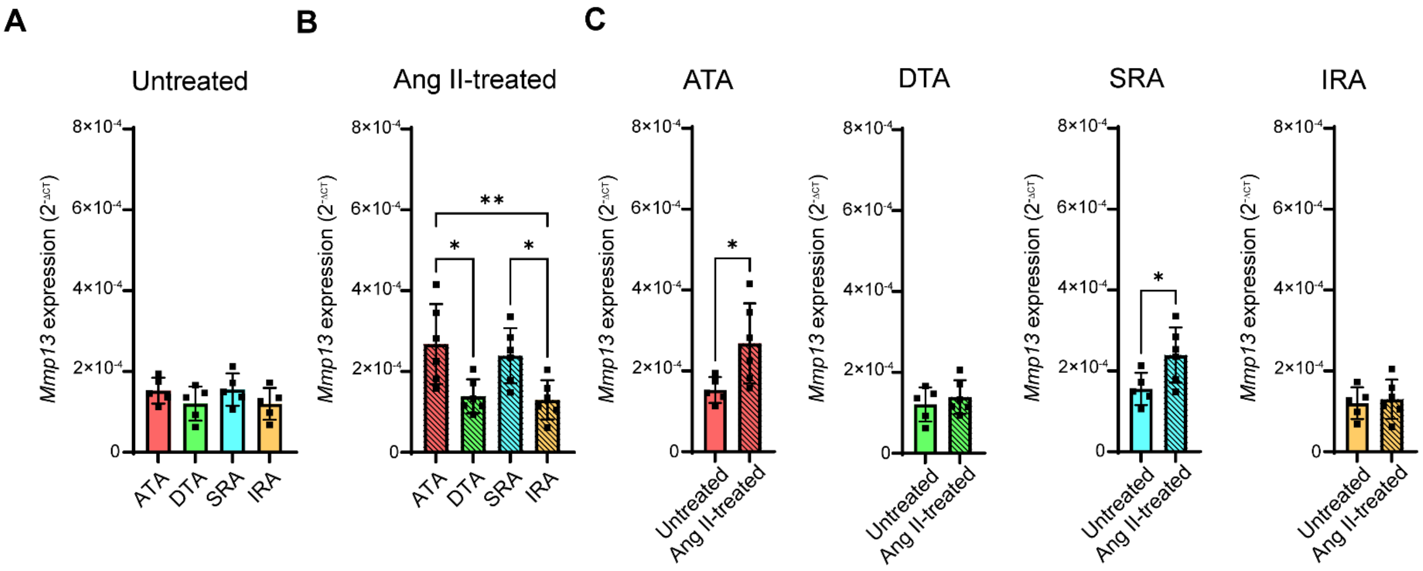


**Supplemental Figure11.** *Mmp13* expression in different segments of the aorta of *Apoe*$^{-/-}$ mice. (12-17 weeks). **A, B.** *Gapdh*-normalized *Mmp13* expression in different aorta segments from untreated *Apoe*$^{-/-}$ mice (**A**), and *Apoe*$^{-/-}$ mice treated with angiotensin II for 3 days (**B**). **C.** Comparison of *Mmp13* expression between untreated and angiotensin II-treated mice within each segment of the aorta. *:*P* < 0.05, **: *P* < 0.01. RFU: Relative fluorescence units, ATA: Ascending thoracic aorta, DTA: Descending thoracic aorta, SRA: suprarenal abdominal aorta, IRA: Infrarenal abdominal aorta.

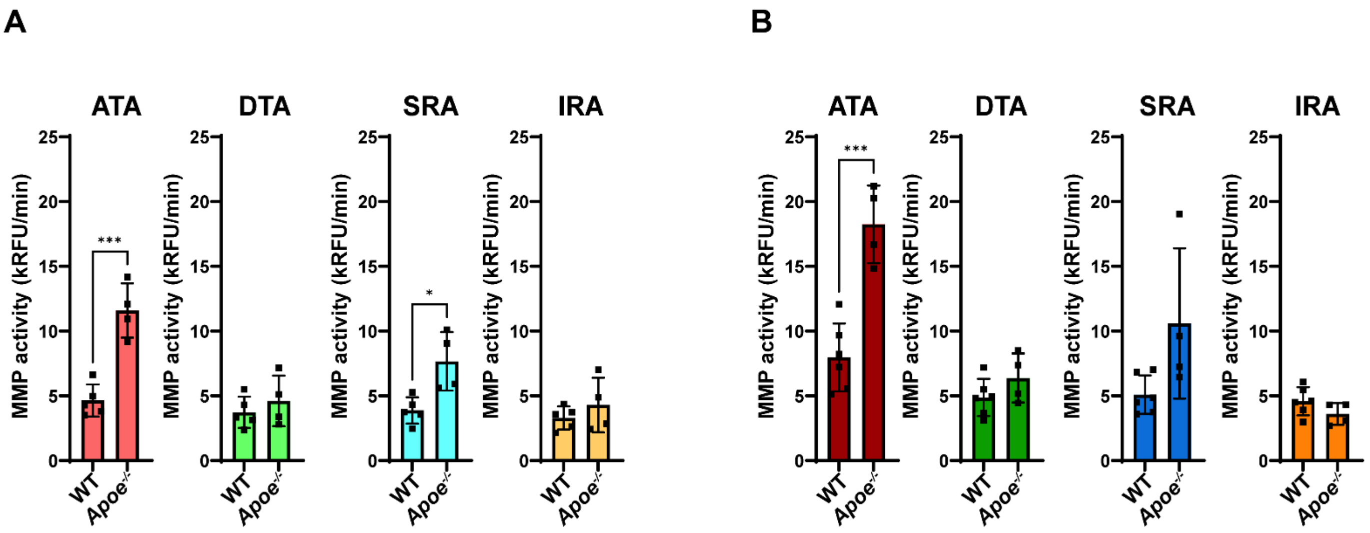


**Supplemental Figure 12.** Comparison of MMP activity in different segments of the aorta of young (8-10 week, **A**) and old (>51 weeks, **B**) wild type and *Apoe*$^{-/-}$ mice. *: $P < 0.05$, ***: $P < 0.001$. RFU: Relative fluorescence units, ATA: Ascending thoracic aorta, DTA: Descending thoracic aorta, SRA: suprarenal abdominal aorta, IRA: Infrarenal abdominal aorta.
